# Transmission mode is associated with environment type and taxa across bacteria-eukaryote symbioses

**DOI:** 10.1101/412882

**Authors:** Shelbi L Russell

## Abstract

Symbiotic associations between bacteria and eukaryotes exhibit a range of transmission strategies. Anecdotal observations suggest that symbionts of terrestrial hosts tend to be strictly vertically inherited through host tissues, whereas symbionts of marine hosts tend towards horizontal transmission. Aside from a few cursory investigations, the rates and distributions of transmission modes have not been investigated in depth across associations, despite the consequences for symbiont and host evolution. To address this empirically and estimate the rate of each mode, I compiled data from the literature and for associations between transmission mode and the environment, transmission route, symbiont function, and taxa involved in the symbiosis. Of the 441 analyzed symbioses, 50.1% were strictly vertically transmitted, 34.0% exhibited some form of mixed mode transmission, and 15.9% were strictly horizontally transmitted. Binning symbioses by their environment revealed a significant skew towards vertical transmission on land and horizontal transmission in aquatic environments, with mixed modes abundant in both. However, host and symbiont taxa were not randomly distributed, which prevented the full separation of these variables. Overall, the data suggest that many symbiotic taxa are capable of horizontal transmission and barriers exist that reduce the rate of these events. Thus, both the environment type and host/symbiont taxa combined influence symbiont transmission mode evolution.

**One sentence summary:** Through an intensive analysis of the literature on symbiosis transmission modes, estimated rates for each transmission mode were calculated, revealing bias for horizontal transmission in the ocean and vertical transmission on land.

## Introduction

An enormous diversity of eukaryotes host bacterial symbionts for a range of functions, from nutrition to defense. In many cases, these associations have allowed both bacteria and hosts to colonize and adapt to radically new niches compared to their ancestral lifestyles. In the nutritional symbioses of sternorrhynchan and auchenorrhynchan insects (Hemiptera), bacterial symbionts allow their hosts to live on diets based solely on phloem and xylem, respectively, and have likely enabled the great radiations of diversity in these groups (Sudakaran *et al.* 2017). As an even more striking example, symbiosis with chemosynthetic sulfur-oxidizing bacteria has evolved multiple times and enabled several marine invertebrate taxa to colonize reducing environments such as the hydrothermal vents. These habitats provided no nutritional or energetic content to the host ancestors, which lacked symbiotic organisms containing the genes for sulfide oxidation and carbon fixation (Stewart *et al.* 2005), underscoring the necessity of these associations for the host’s niche. Defensive symbionts often provide contextual benefit to their hosts, for example, by providing antibacterial defense during embryogenesis (Kaltenpoth *et al.* 2010 and Flórez *et al.* 2017) or protection against parasitoids in environments where the threat is present (Oliver *et al.* 2010). While some symbionts appear to perform multiple functions (*e.g., Steinernema carpocapsae* of pathogenic nematodes which are involved in host nutrition and development (Goodrich-Blair *et al.* 2007)), the vast majority would appear to specialize in a task. Alternatively, manipulative associations do exist in which symbionts do not provide a benefit, but tie themselves to host reproduction and so have to act within host interests. Many of the associations between *Wolbachia* and its invertebrate hosts fall into this category. These bacteria generally achieve high infection rates through efficient vertical transmission and reproductive manipulations (Serbus *et al.* 2007; Werren *et al.* 2008; Russell *et al.* 2018). In cases such as filarial nematodes (Landmann *et al.* 2011; Foray *et al.* 2018) and *Asobara* wasps (Dedeine *et al.* 2005), *Wolbachia* have evolved to control host development, making these associations appear like mutualisms. However, in these situations *Wolbachia* has inserted themselves into core host functions, including cell differentiation and proliferation, making this more of an addictive association (Sullivan 2017).

Regardless of the function, symbionts and hosts must find each other every generation via navigation through host tissues or the environment in order to perpetuate the association over evolutionary time. The routes symbionts take between hosts, or the transmission modes, exist on a gamut between horizontal transmission of symbionts, through an environmental intermediate, and vertical transmission of symbionts via inheritance through host tissues or secretions (described Figure 1A). Mixed-mode transmission incorporates some rate of horizontal and vertical transmission to achieve reliable host association (Bright and Bulgheresi 2010). Horizontal transmission via uptake of bacteria from the environment, perhaps migrating from other contemporary hosts, is most certainly the ancestral mode for all symbioses because all known symbionts, even mitochondria and chloroplast organelles, have free-living ancestors (Gray *et al.* 1999). Many associations have kept with this strategy (*e.g.*, hydrothermal vent tubeworm symbionts (Nussbaumer *et al.* 2006)), however, a great many others have evolved elaborate mechanisms of symbiont inheritance ranging from egg-smeared secretions (*e.g.*, stink bugs (Hosokawa *et al.* 2013)) to incorporation of symbionts in developing oocytes (*e.g.*, vesicomyid bivalves (Ikuta *et al.* 2016)) or embryos (*e.g.*, parthenogenetic aphids (Braendle *et al.* 2003)). In a larger number of associations, there is data that supports both vertical and horizontal transmission, suggesting that mixed-mode transmission is quite common. Mixed modes can arise via either the occurrence of horizontal events in a faithfully vertically transmitted association (*e.g.*, solemyid bivalves (Russell *et al.* 2017) and aphid facultative symbionts (Vorburger *et al.* 2017)) or one in which vertical transmission fails at a high rate (*e.g.*, chinch bug vertical transmission fails 70% of the time (Itoh *et al.* 2014)), placing more importance on the horizontal transmission strategy (described in Figure 1A).

**Figure 1.**
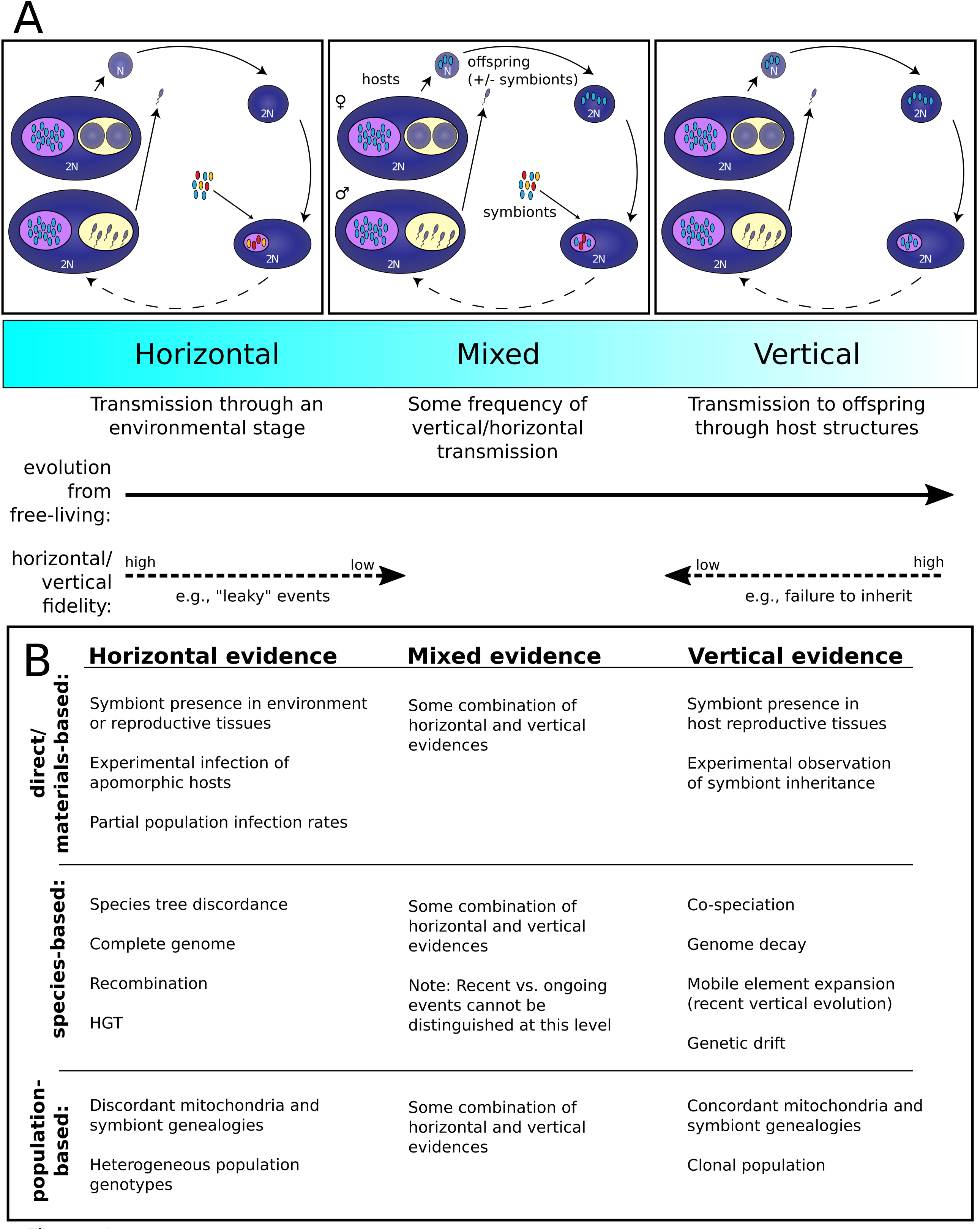
Transmission mode spectrum definitions and influential parameters. A) Symbionts evolve from free-living ancestors, and are thus initially horizontally transmitted through an intermediate environmental stage. Some associations evolve vertical transmission through host tissues, either obligately or in conjunction with some amount of horizontal transmission. B) Criteria for binning symbioses into each transmission mode based on data from the literature (columns labeled in colored bar A).

Recent genomic population-level data has rekindled interest in mixed-mode transmission, as rare horizontal events substantially impact bacterial symbiont genome evolution (Russell *et al.* 2017), even if they are not informative predictors of host-symbiont cooperation (Fisher *et al.* 2017 and references therein). In strictly vertically transmitted associations such as those between sap-feeding insects such as aphids and spittlebugs and their symbionts *Buchnera* and *Sulcia*, respectively, host restriction and genetic drift over millions of years has resulted in severe genome erosion (Moran and Bennett 2014), with many symbiont lineages lacking even the genes needed for the association, requiring the acquisition of a secondary symbiont with complementary functions (*e.g.*, spittlebugs (Koga and Moran 2014)). While a vast number of associations fit this model, many of which are hemipteran-associated, some associations exhibit vertical transmission without such severe genome reduction (*e.g.*, Woyke *et al.* 2006; Newton *et al.* 2007; Dmytrenko *et al.* 2014). Given that many of these associations occur in marine habitats, and vertical transmission has been reported to be rare in this environment (Normark and Ross 2014), habitat type may shape the cost/benefit tradeoffs for transmission modes. Intuitively, this is reasonable as water is a much easier medium for a bacterium to navigate through than air. Thus, in this work I investigated whether there is indeed a non-random distribution of symbiont transmission modes among environment types.

## Materials and Methods

### Literature search

To assess the rate of mixed mode transmission among bacteria-eukaryote symbioses, I obtained references from either searches with Google Scholar, using keywords such as “transmission mode”, “inheritance”, “infection”, “colonization”, “transovarial transmission”, “symbiont uptake”, “bacterial/symbiont phagocytosis”, “bacterial/symbiont endocytosis”, etc., a Zotero database of papers on the subject of symbiont transmission I had been collecting for years, or the reference sections of these papers. This search produced over 250 journal articles containing transmission mode data of either a direct or indirect nature for 441 symbiont-host associations were analyzed (Figure 1B and Table S1, supporting information). While many more papers than these 250 exist that characterize symbioses in general, and include associations not presented in our analysis, only the papers and associations that had some amount of data on transmission mode were included.

### Data collection

#### Definition of bacteria-eukaryote symbioses and transmission modes

The associations included in this analysis were those between an identified bacterial symbiont strain and a eukaryotic host that are either mutually beneficial (*i.e.*, mutualisms), or are beneficial for one partner and commensal for the other. This second category of nearly-mutualistic associations is important to consider in regards to transmission, as different evolutionary dynamics may influence them. Symbionts in this category of association often manipulate host reproduction via a strategy such as cytoplasmic incompatibility, feminization, parthenogenesis, or male killing (Werren *et al.* 2008; Ma *et al.* 2014; Correa and Ballard 2016; Larracuente and Meller 2016), enabling maintenance of the association despite not providing a benefit to the host. I assigned the function “manipulative” to symbioses in this category because this is the predominant term used in the literature (*e.g.,* Werren *et al.* 2008; Correa and Ballard 2016). I also tallied the mutualisms according to the functions provided to the host, as this is a common approach for binning symbioses (*e.g.,* Toft and Andersson 2010; Douglas 2016; Sudakaran *et al.* 2017) and provides insight into host-symbiont interaction. Associations were binned into either “nutrition”, such as chemosynthetic or photosynthetic carbon fixation, nitrogen fixation, or amino acid synthesis, “defense”, such as viral or parasitoid protection, antibiotics, or bioluminescence, “multiple function” for the associations that perform both a nutrition and a defense role (*e.g., Serratia symbiotica* provides both defensive and nutrition functions in aphids (Oliver *et al.* 2010; Burke and Moran 2011)), “unknown” for mutualisms that have not been functionally characterized, or “manipulative” as described previously. In addition to symbiosis function, I collected data on vertical transmission routes and environment types (see below), as these factors may have an influence on transmission modes. Microbiome-type associations were not included because the taxa involved are rarely identified at a fine enough scale to permit comparisons among datasets, prohibiting synthesis of this literature. Lastly, it should be noted that both intracellular and extracellular symbioses were included in this analysis, as it has been shown that the cellular environment does not prevent the evolution of traits thought to to be unique to an intracellular lifestyle, such as genome reduction (Nicks and Rahn-Lee 2017; Salem *et al*. 2017).

Intuitively, transmission modes exist on a continuum from strict horizontal transmission at one end to strict vertical transmission at the other, with mixed modes describing the strategies in between (Figure 1A). While it would be ideal to treat transmission mode as a continuous character, reporting on mixed modes is still in its infancy and precludes more quantitative analyses. For example, few papers do more than show symbiont presence in host reproductive tissues or infer a multi-species phylogeny, making precise estimates of mixed mode rates impossible to obtain at this time. Thus, for the purposes of this work, transmission mode is defined as a discrete trait with three values, “horizontal”, “mixed”, and “vertical”.

For vertical transmission, routes between host generations can either exclusively involve intra-host transfers or can involve some external intermediate, which could influence the rate of horizontal transmission in mixed modes. For example, while parthenogenetic aphids brood their young, they transfer their *Buchnera* symbionts from adult bacteriocytes to embryonic cells (Koga *et al.* 2012), exemplifying what I’ve defined as “internal vertical transmission”. In contrast, and demonstrating “external vertical transmission”, pentatomid stinkbugs smear a symbiont-containing secretion on their eggs after they lay them (Hosokawa *et al.* 2013).

Lastly, the environment type, marine, terrestrial, or freshwater, was noted for each association, as the medium may impact the evolution of transmission modes (Normark and Ross 2014). As only two freshwater associations were found with transmission data and the medium, water versus air, is thought to be the critical factor, marine and freshwater tallies were pooled as “aquatic” associations, and the remainder “terrestrial”.

#### Transmission mode frequency estimation

Few papers directly report mixed transmission modes, in large part because appreciation for this intermediate mode has grown recently and the assessment of mixed modes involves multiple data/experiment types. Far more common are papers that state results departing from expectations for strict vertical transmission (*e.g.*, genealogical or phylogenetic discordance). Alternatively, pairs of papers show direct evidence of vertical or horizontal transmission (*e.g.*, through host tissues or environmental substrates, respectively) in one and genetic evidence of departures from strict vertical/horizontal transmission in another.

To arrive at an estimate for the rate of horizontal, vertical, and mixed mode transmission among symbiotic associations, these miscellaneous and partial data from the literature had to be collected and synthesized. Thus, I inferred mixed modes from a combination of direct evidence, such as symbiont presence in germline tissue, mixed infections, or experimental horizontal transfers, and indirect evidence, such as a lack of co-speciation, evidence of recombination or HGT, or a lack of genealogical discordance (for studies on populations). It should be noted that in the definition of mixed transmission used here, no distinction is made between horizontal and paternal transmission, as this was generally not tested for in the literature and is hard to distinguish otherwise. However, rates of paternal transmission are generally low and are often mediated via host-to-host contact (essentially a horizontal transmission process), as sperm are too small to house many bacterial symbionts (Peccoud et al. 2014; De Vooght *et al.* 2015). In addition, no distinction was made between ongoing horizontal transmission and the recent evolution of strict vertical transmission, as complete taxonomic sampling and a time-calibrated phylogeny is needed to state this with confidence (as in Lefevre *et al.* 2004), which is not available for most associations. Lastly, only bacterial symbionts were considered, however, replacements with yeast symbionts have occurred in the planthoppers (Bennett and Moran 2015), and cnidarian zooxanthellae exhibit a range of transmission modes (Quigley *et al.* 2018).

Transmission evidence was binned into one of three categories: A) direct/material-based evidence, B) indirect species-level evidence, and C) indirect population-level evidence (see Figure 1B). Direct, or materials-based, evidence included any observation of symbionts in the reproductive tissues (vertical transmission) or at high abundance in the environment (horizontal transmission). In a few cases, symbiont experimental infection studies had been performed in the lab, providing the most direct of data supporting horizontal transmission (*e.g.*, Gonella *et al.* 2012). Indirect data were categorized at two levels: species-level evidence and population-level evidence. Species-level data was by-far the most common, as a great number of studies over the past few decades have PCR-amplified and Sanger sequenced symbiont markers, often the 16S ribosomal RNA, and host markers, often mitochondrial loci such as COI, to infer the host-symbiont co-evolutionary relationship. However, the advances in sequencing technologies over the past 15 years have permitted many symbiont genomes to be sequenced, providing data on whether the genome exhibits evidence of strict vertical transmission, *i.e.*, genome size erosion, gene loss, AT-bias, accelerated substitution rate, etc. (Andersson and Kurland 1998), or resembles that of a free-living bacterium. Recently evolved strict vertical transmission is hallmarked by extensive mobile element expansions (Plague *et al*. 2007; Newton and Bordenstein 2011). Additionally, these studies have also revealed evidence of rampant recombination in some symbiont genomes (Baldo *et al*. 2005; Russell *et al.* 2017; Russell and Cavanaugh 2017), which requires some rate of horizontal transmission to be observable. At the population level, evidence of vertical transmission includes a clonal intra-host population and mitochondria-symbiont genealogical concordance. While genealogical discordance is too sensitive of a measure to distinguish degrees of horizontal transmission, being disrupted with even exceedingly low rates (Brandvain *et al.* 2011), genetic evidence of mixed populations in host tissues is strong evidence that horizontal transmission is ongoing at a high rate (Russell and Cavanaugh 2017). Transmission evidence levels and results are presented in the “evidence for mode” column of Table S1, supporting information, and are indicated as follows: direct evidence in support of vertical transmission equals 1 or horizontal transmission equals 2, species-level evidence in support of vertical transmission equals 3 or horizontal transmission equals 4, and population-level evidence in support of vertical transmission equals 5 or horizontal transmission equals 6.

The patchiness of the available data on symbiont transmission made it impossible to use the same criteria to call transmission modes across all associations. Thus, conservative calls were made from what data could be obtained. For example, if only tissue/environmental-based data existed in support for vertical/horizontal transmission, and no genetic data was available on the fidelity of that mode, then the strict vertical/horizontal transmission mode was assigned. This conservative approach likely produced an underestimate of the frequency of mixed modes among symbioses.

### Statistical methods

Data for the 441 symbioses was sorted, tallied by category (transmission mode, environment, (vertical) transmission route, and function), and output in contingency tables with a custom perl script. These tables were read into R for plotting and statistical testing. Fisher’s exact test was used on these nominal data to determine whether the relative transmission mode rates are independent of environment type and whether vertical transmission route, external or internal, is independent of transmission mode. Pearson’s chi-squared test was used to test for an impact of symbiosis function on transmission mode. Associations between host/symbiont taxa and the tallied variables were plotted using the heatmap.2 (ggplot) heatmap plotting function in R.

Permutation tests were performed on the data to determine whether enrichments for transmission modes in the different categories were significant above noise/bias in the dataset. To do this, the data table in Table S2, supporting information was read into a custom perl script, and the columns were shuffled, randomizing the data for each symbiosis. The categories described above were tallied for these permuted data, and recorded. This was then repeated for a total of 1000 times to determine how many times the empirical data’s value was above/below the randomized data values, and thus the probability the value could have been obtained by chance.

### Sensitivity analyses

Each of the four analyses, 1) transmission rate estimation, 2) correlations with environment type, 3) mixed mode correlation with external vertical transmission routes, and 4) interplay between function and transmission mode, were repeated on subsets of the data as described below.

### Robustness to data availability

To maximize the amount of data/number of associations included in the analysis, I included associations that had at least one form of evidence among materials-based, species-based, and population-based evidence. However, this could bias the results, if lack of data drives the dominance of one more or another. For example, there may be a skew towards vertical transmission when only host tissues have been examined. To account for this, I repeated the analyses two more times, once on the subsets of associations that had at least two levels of evidence, and again on the subset that had all three. Subsampling by evidence level reduced the dataset to 324 associations with a minimum of two levels and 102 with all three.

### Robustness to taxon selection

To assess the impact dense taxonomic sampling (esp. insects) had on the results presented in Figure 2, I subsampled two host and symbiont taxonomic levels, one at a high-level (class) and one at a lower level (family). After finding the minimum number of unique symbioses at each of these taxonomic levels, I randomized and resampled the data to only include one of each in a subsampled dataset with a custom perl script, which I then analyzed as described above. This reduced the dataset to 7-10 symbioses when classes/phyla can only be represented once and 36-43 symbioses when families can only be represented once, depending on the host/symbiont taxonomic combinations selected. This was repeated 1000 times for each high-level and low-level dataset.

**Figure 2.**
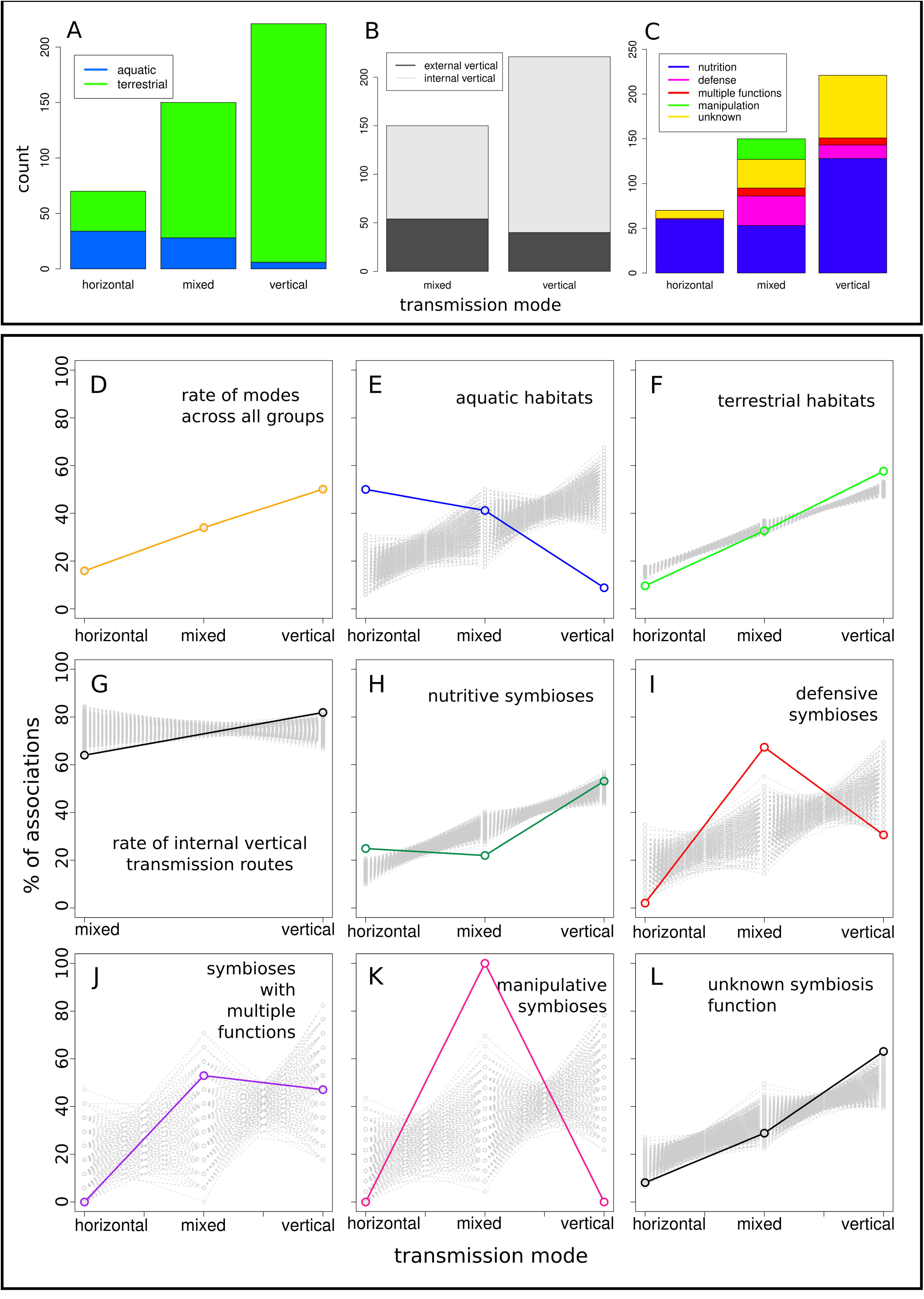
Transmission mode distribution (D) across A,E,F) environment types, B,G) transmission routes, and C,H-L) symbiosis functional types. Transmission mode values for the total dataset are plotted in color with solid lines connecting the points in D-L, whereas values for the randomly permuted datasets are plotted in grey with dashed lines connecting the points. Color keys for the bar plots are indicated in each panel in A-C, and correspond with their respective full dataset colors in D-L, as well as in Figures 3, 4, and S1.

### Robustness to unequal sampling or biodiversity distribution

Similarly to the dense taxonomic sampling situation, the accumulated dataset is enriched in terrestrial associations, which could bias results. To control for this, I randomly subsampled the terrestrial symbioses to match the sample size of aquatic symbioses (n = 68) and calculated transmission mode rates. This was repeated 1000 times.

Resulting datafiles were plotted in R. The scripts used to perform these analyses can be downloaded from https://github.com/shelbirussell/MixedModeAnalysis_Russell2018.

## Results

Tallying transmission mode counts over all 441 symbiotic associations shows that 15.9% (70/441) of symbioses considered here are strictly horizontally transmitted, 34.0% (150/441) have some degree of mixed transmission, and 50.1% (221/441) are strictly vertically transmitted (Figure 2A-D). While these numbers would indicate that there is some overall preference for associations to adopt vertical transmission relative to horizontal transmission, it should be pointed out that the sampling distribution is highly skewed towards terrestrial associations (373 terrestrial versus 66 aquatic; Figure 4A), which may either be due to sampling bias or a true excess of terrestrial species relative to marine and freshwater species in terms of biodiversity (Figure 4B; Grosberg *et al.* 2012, Costello and Chaudhary 2017). If the symbioses are randomized and subsampled to equalize sample counts across taxa and environment type, this bias disappears, and the modes become approximately equally distributed (Figure 3C and 4C). Regarding how the quality of the data affects these results, the values remained relatively consistent when two levels of evidence were required for inclusion in the dataset (Figure S1, supporting information). However, when all three levels were required, the estimates for mixed modes increasing in almost all cases, suggesting that the full dataset underestimates the rate of mixed modes.

**Figure 3.**
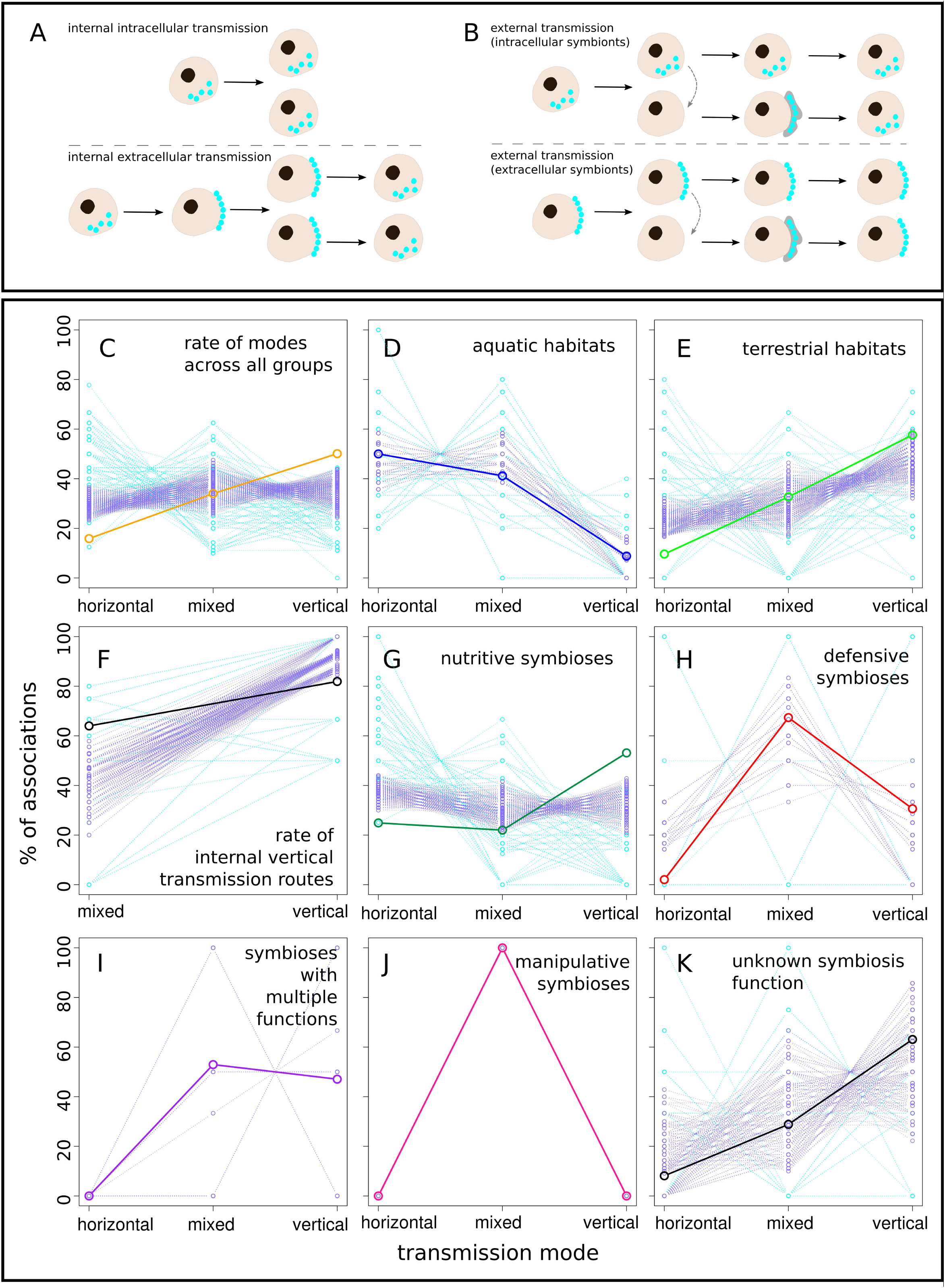
Transmission mode distribution (C) across D,E) environment types, A,B,F) transmission routes, and G-K) symbiosis functional types compared to randomized datasets consisting of subsampled taxa. A,B) Illustrations of the defined A) internal and B) external vertical transmission routes, with transfer steps shown for intracellular (top) and extracellular (bottom) associations. Transmission mode values for the total dataset are plotted in color with solid lines connecting the points in D-L, and correspond to the colors used in Figures 2, 4, and S1. Values for the randomly subsampled high taxonomic level (blue) and low taxonomic level (purple) datasets are plotted with dashed lines connecting the points.

**Figure 4.**
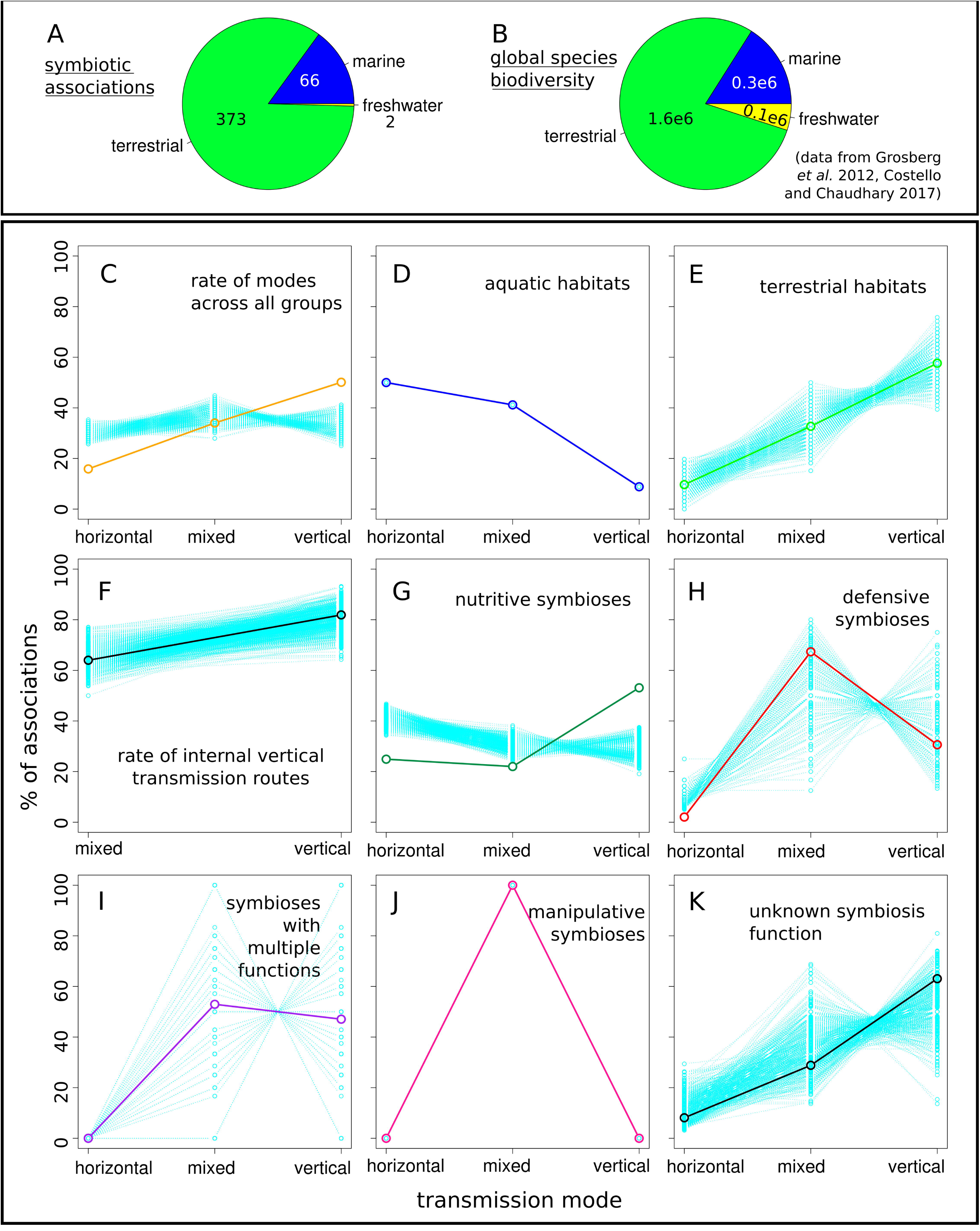
Transmission mode distribution (C) across A,B,D,E) environment types, F) transmission routes, and G-K) symbiosis functional types compared to randomized datasets consisting of subsampled terrestrial samples. Distribution of A) symbiotic associations and B) global species diversity between marine, terrestrial, and aquatic environments. Transmission mode values for the total dataset are plotted in color with solid lines connecting the points in D-L, and correspond to the colors used in Figures 2, 3, and S1. Values for the randomly subsampled environmental datasets (in which the number of terrestrial samples = aquatic samples) are plotted with dashed teal lines connecting the points.

Compiling these data and binning by variables such as transmission route, environment type, symbiont function, and host/symbiont taxon revealed that symbioses are indeed non-randomly distributed (Figure 2). However, it should be noted that there is no way to test this question appropriately with the full dataset because the distribution of taxa is itself non-random, making phylogenetically independent contrasts impossible. For example, neither bivalves nor chemosynthetic ecosystems occur on land and insects make very limited ventures into aquatic environments of any sort (*e.g.*, reed beetles (Kölsch and Pedersen 2010) and the seal louse (Boyd *et al.* 2016)). Therefore, to control for the impact of repeated sampling of close relatives, I performed 1000 iterations of taxonomic subsampling (Figure 3). To control for the excess of terrestrial associations reported (Figure 3A), I randomly subsampled the terrestrial associations to equal the sample size of aquatic associations 1000 times (Figure 4). These results show that transmission mode is indeed correlated with variables such as environment type.

### Transmission modes are non-randomly distributed among environments, routes, and functions

These data show that terrestrial environments are indeed enriched for vertical transmission and aquatic environments are enriched for horizontal transmission (Figure 2A,E,F), consistent with the observations of Normark and Ross (2014). Comparing the distribution of these three modes between terrestrial and aquatic environment types reveals a highly skewed distribution (Fisher’s Exact Test p < 2.2e-16; Figure 2A and Table S2, supporting information). Terrestrial environments are greatly enriched for strict vertical transmission, with 57.6% (215/373) of terrestrial symbioses exhibiting this mode. Furthermore, as 97.2% (215/221) of vertically transmitted associations occur on land (Figure 2A), this mode appears to be virtually restricted to this environment. Compared to the permuted dataset, the enrichment in horizontal transmission and depletion of vertical transmission in aquatic environments are highly significant (permutation test p < 0.001; see the data’s negative slope vs the permutations’ positive slopes across the transmission categories in Figure 2E). Similarly, the terrestrial environment is significantly depleted for horizontal transmission and enriched for vertical transmission, with no permuted value below/above the empirical values, respectively (permutation test p < 0.001). Mixed transmission is also skewed more towards terrestrial associations (81.3%, 122/150). However, compared to the permuted values, there is depletion or enrichment for mixed modes in either environment (permutation test p = 0.17 and 0.08 in aquatic and terrestrial habitats, respectively).

When these data are randomly subsampled to only include non-redundant taxa or equal numbers from each environment type, these trends persist, indicating that the interactions between variables are large-scale impacts and are not driven by a small number of associations. The signal in aquatic habitats appears to be driven by a diversity of taxa, as the transmission mode rates for the full dataset are well within the values for the permuted taxa dataset (Figure 3D). As aquatic environments were the lower-abundance environment type, these values were not subsampled and so these values overlap with the full dataset in Figure 4D. Lying closer to the extremes, the terrestrial environment transmission mode rates were barely within the subsampled high-level taxa values (Figure 3E), with 7/1000 subsampled values lying below the terrestrial horizontal transmission mode rate and 102/1000 above the vertical transmission mode rate. Furthermore, none of the subsampled low-level taxa values were below the terrestrial horizontal transmission mode rate and only 8/1000 lay above the vertical transmission mode rate, suggesting that only a few taxa drive this aspect of the signal. When terrestrial associations are randomly subsampled, these values overlap the full dataset values (Figure 4E), indicating that the abundance of terrestrial symbioses does not contribute to the elevated vertical or depressed horizontal transmission mode signal.

### External modes of vertical transmission may predispose associations to mixed mode transmission

Next, the counts of external versus internal vertical transmission routes in each transmission mode were compared to assess whether transmission route may contribute to the evolution of mixed modes (Figure 2B). The logic behind this idea is that although, strictly speaking, externally vertically transmitted symbionts are inherited, and often with very high fidelity (Newton *et al.* 2015; Vorburger *et al*. 2017), the external route of transmission offers opportunities for symbiont mixing that do not occur if symbionts never leave the host body (Figure 3A,B; Salem *et al.* 2015).

Comparing transmission routes for symbiont inheritance in mixed and vertical modes of transmission revealed a significant deficit of internal transmission routes in mixed modes relative to vertical transmission (Fisher’s Exact Test p = 1.5e-4; Figure 2B). 36.0% (54/150) of associations with mixed modes utilize some form of external vertical transmission. This strategy ranges from secretions applied to eggs, which occurs in stink bugs (Hosokawa *et al.* 2013), beewolves (Kaltenpoth*et al.* 2006), reed beetles (Kölsch and Pedersen 2010), earthworms (Paz *et al.* 2017), and chemosynthetic oligochaete worms (Giere 2006), to elaborate symbiont-containing capsules laid with eggs, which occur in plataspid stink bug species (Hosokawa *et al.* 2005). In contrast, only 18.1% (40/221) of strictly vertically transmitted associations exhibit an external route of transmission. The permuted values were greater than the mixed mode internal transmission rate and less than the vertical mode internal transmission rate (permutation test p = 0.002 and 0.001; Figure 2G), further supporting a non-random distribution of transmission routes between transmission modes. However, while, the internal route proportion values from the full dataset are well within the subsampled high-level taxa (Figure 3F, light blue) and environment (Figure 4F) datasets, these values were outside the subsampled low-level taxa values (Figure 3F, purple), indicating that some symbiont or host families may drive this signal.

### Transmission modes are non-randomly distributed among symbiosis functional types

Next, I binned symbioses by one of six functional types and evaluated their distribution among modes(Figure 2C,H-L), revealing a highly non-random distribution (Pearson’s Chi-squared test p < 2.2e-16). To explore how the modes are distributed in each functional type, we first compared them to randomly permuted distributions of values (Figure 2H-L).

In symbioses that are based on nutrition functions, horizontal transmission is elevated and mixed modes are depleted relative to permuted values (permutation test p < 0.001), whereas vertical transmission is not significantly different from the randomized distribution (permutation test p = 0.05). Elevated horizontal transmission is likely driven predominantly by the chemosynthetic portion of the nutritional symbioses, as horizontal transmission is enriched for chemosynthetic associations, and vice versa. However, as all known chemosynthetic symbioses are marine (47/47), this association cannot be disentangled from that between horizontal transmission and the aquatic environments in general (Figure 2A,E). Subsampled high and low-level taxon and environment distributions for nutritional symbioses were also enriched for horizontal transmission and depleted for vertical transmission relative to the total dataset (Figure 3G and 4G). In the case of horizontal transmission, the rate of which was already significantly higher than the permuted values (Figure 2H), such that the subsampled values exhibit even higher rates of horizontal transmission than the full dataset, indicating that a good range of families contribute to the horizontal transmission enrichment in nutritional symbioses.

Defense, multiple function, and manipulative associations were highly depleted for horizontal transmission, with 1/49, 0/17, 0/23 associations exhibiting exclusive horizontal transmission, respectively. Compared to permuted datasets, defensive and manipulative symbioses were depleted in horizontal (permutation test p < 0.005 and 0.01, respectively) and vertical modes (permutation test p < 0.001), whereas they were enriched for mixed modes of transmission (permutation test p < 0.001), distinguishing these factors from the terrestrial environment factor (Figure 2I). Symbioses with multiple functions were depleted in horizontal transmission relative to permuted values (permutation test p < 0.03), but had expected rates of mixed and vertical modes (Figure 2J), exhibiting an intermediate pattern to the nutritional and defensive functional components that contribute to this functional category. However, these functions are predominantly seen in terrestrial environments (45/49, 17/17, and 21/23 in defensive, multiple function, and manipulative associations, respectively), and so horizontal transmission may be limited by that factor.

The associations in the unknown bin exhibited enriched vertical transmission (permutation test p= 0.01), depleted horizontal transmission (permutation test p < 0.001), whereas mixed modes were no different from the permutations (permutation test p = 0.09). These values are similar to those for terrestrial and/or nutritional symbioses (Figure 2F,H).

Subsampled taxa were largely consistent with the full dataset values for defensive, manipulative, and unknown function symbioses (Figure 3H,J,K), and the permuted environmental datasets were largely overlapping as well (Figure 4H,J,K). However, the number of subsampled sets was small for defensive and manipulative symbioses, likely because few symbiont taxa exhibit these phenotypes, and the subsampled high-level taxon datasets produced highly variable results for defensive (Figure 3H), limiting the conclusions that can be drawn from these data. While the subsampled environment dataset values overlapped with those for the full dataset for manipulative symbioses (Figure 4I), both the low and high taxonomic level subsampled data sets produced variable results and were of low count, probably also due to few symbiont taxa exhibiting this phenotype.

### Phylogenetic effect: Transmission modes are non-randomly distributed among symbiont and host taxa

While the correlations between transmission mode and these biological and environmental variables are compelling, and it is clear that many symbioses contribute to these trends, it is possible groups of host or symbiont taxa that covary with the variables control transmission mode. Unfortunately this cannot be tested directly, as the naturally constructed phylogenetically-independent contrasts do not exist (*e.g.*, chemosynthetic symbioses on land). However, the relative correlations between each of the components and the taxa can be examined and compared to qualitatively answer this question.

Analysis of the clustering among symbiosis traits and taxa revealed that the factors exhibited strong covariation with one another, confirming that they are not randomly distributed. As shown in Figures S2 and S3, supporting information, both taxonomic divisions of symbionts and hosts showed strong patterns of mutual-exclusivity and specificity for factor combinations (dendrograms on x-axes in Figure S2 and S3, supporting information). The unique combination of traits specific to insect symbioses, terrestrial [environment], nutrition [function], vertical [mode], and internal route, were clustered to the exclusion of the other traits, and were specific to insects and insect phyla as well as their symbiont clades, *e.g.*, Enterobacteriaceae. Aspects of the marine chemosynthetic symbioses cluster, however, they do not form as cohesive of a group when chemosynthesis is included in the broad category of nutrition functions as they do when chemosynthesis is considered separately. The other groupings were less well-resolved, suggesting more sharing of these other traits among taxa. Mixed modes is itself one of these traits, as it is fairly well-distributed across taxa, with conspicuous absence in only a few groups. Defense, being a fairly broadly-defined trait that ranges from the production of bioluminescent light in squid (Nyholm and McFall-Ngai 2004) to antibiotic production in beewolves (Kaltenpoth *et al.* 2010) and beetles (Flórez *et al.* 2017), to cuticle hardness in beetles (Anbutsu *et al.* 2017), was also distributed across taxa. Additionally, external vertical transmission routes, such as egg-smearing, were also highly dispersed, but often grouped with mixed modes.

Regarding taxon clustering (y-axis dendrograms in Figure S2 and S3, supporting information), the topology of these clustering trees had very short terminal branches nested within one another, with few deep branches describing broad-scale groupings of taxa. This suggests that while the factors analyzed here are non-randomly distributed among taxa, the taxa themselves consist of unique combinations of these traits, with few forming “guilds” that all exhibit similar properties.

## Discussion

Every symbiotic association is posed with the same challenge: How do symbionts establish contact with host offspring during host reproduction? A range of solutions have been adopted across the diversity of symbioses, but little is known about the factors influencing when each of these solutions evolve. While recent work by Fisher *et al.* 2017 showed that host dependence is correlated with transmission mode and symbiont function, they did not evaluate mixed transmission modes or test for an impact of environment type or transmission route. Thus, a wide scale analysis across the full diversity of bacteria-eukaryote symbioses was warranted. First I sought to quantify the rates of horizontal, mixed, and vertical transmission modes in nature from the literature to date on symbiont transmission. Next, using these data, I tested whether factors such as the environment in which a symbiosis lives, the route symbionts take between host tissues, and the function of the symbiosis have an impact on the distributions of these transmission modes.

Here, I show that the three symbiont transmission mode categories, horizontal, mixed, and vertical are unequally distributed across environment types, transmission routes, and some functional types, indicating that these factors either pose some constraint or pressure favoring more or less horizontal transmission. These results are robust, as many of the trends persist when host/symbiont taxa and environments are subsampled. The most striking of these trends is the enrichment of horizontal transmission and depletion of vertical transmission in aquatic environments (Figure 2A,E,F), the rate of internal/external transmission routes between mixed and vertical modes of transmission (Figure 2B,G), and the enrichment of mixed modes in multiple function and manipulative associations (Figure 2C,I,K). However, it should be kept in mind that phylogeny was unable to be controlled for in these analyses due to the questions being asked (*e.g.*, environment type) and the constraints imposed by the distribution of symbioses in nature, as chemosynthetic bivalves do not occur on land and sap-feeding insects do not occur in the ocean. Therefore, while some host and symbiont taxa are specific to the variables tested (Figure S2, supporting information) and may drive the observed associations, the randomized taxon subsampling analysis supports that the broadscale patterns observed are not specific to the taxa sampled (Figure 3C-K).

The strong correlation between terrestrial symbioses and strict vertical transmission, and marine symbioses and horizontal transmission, as either strict horizontal or mixed mode transmission, may simply be an artifact of the medium in which these organisms live: water versus air (Normark and Ross 2014). With desiccation and osmolarity not a problem, symbionts can travel through aqueous environments to new hosts if the opportunity arises and they are capable of surviving outside of the host cell. Not much fluid may be needed, as all nitrogen-fixing rhizobia root bacteria are horizontally transmitted through soil pore water (Postma and van Veen 1990). The marine environment to which most of the investigated aquatic symbioses belonged (66/68) likely further enables host-to-host transfer events via its salt and nutrient content (Rozen and Belken 2001).

The dependence of horizontal transmission on environmental opportunity is supported by finding that external transmission routes are more common in mixed modes and literature on the intracellular alphaproteobacterial symbiont *Wolbachia.* First, if transmission mode was fully intrinsic to the host/symbiont taxa and independent of opportunities for alternative strategies, then one would expect there to be no relationship between transmission mode and vertical transmission route. However, finding that mixed modes are enriched for external routes suggests that these strategies, which often involve external applications of symbiont-containing fluids, present opportunities for “contamination” between hosts. In the case of *Wolbachia*, this vertically-transmitted bacterium has been shown to survive for days outside of a host cell in cell culture media (Rasgon *et al.* 2006) and can exit/enter host cells readily in this environment (White *et al.* 2017). Given that horizontal transfer and host-switching events have been inferred to be much rarer in nature than this behavior in culture would suggest, it would appear that *Wolbachia* is restricted from achieving its maximum horizontal transmission rate, possibly by the environment.

The influential role of environmental medium on transmission mode is highlighted by the many terrestrial symbionts functionally capable of horizontal transmission that have been shown to utilize plant fluids and parasitoids as strategies for transfer. Rickettsial symbionts of whiteflies can be transferred through the fluids of the plants the hosts feed on (Li *et al.* 2017). Furthermore, plants may serve as reservoirs for symbionts, facilitating transmission, as some defensive symbionts have been found to also be pathogens of plants (Flórez *et al.* 2017) and the *Erwinia* and *Pantoea* symbionts of stink bugs are also plant pathogens (Frago *et al.* 2012). Transfer between hosts can also be facilitated via infecting a parasitoid wasp, as has been reported for *Wolbachia* (Vavre *et al.* 1999), likely due to its aptitude for surviving in a diversity of arthropod hosts. However, while some symbionts can be transferred through these strategies, it may be a very rare process and is highly dependent on symbiont taxon. For example, it has been shown that different mealybug species with different nested beta/gammaproteobacterial symbionts feeding on the same plant were found to maintain their native symbiont types (Koga *et al.* 2013), despite the gammaproteobacterial symbiont showing evolutionary evidence of host-switching (Gatehouse *et al.* 2012). Thus, it is likely that both the environmental medium and the host/symbiont taxon synergize to determine the specific transmission mode for the symbiosis.

The marine environment appears to represent the most-limiting situation for vertical transmission, as no associations with robust evidence sampling have been found to be strictly vertically transmitted (Stewart *et al.* 2008; Stewart *et al.* 2009; Decker *et al.* 2013; Altamia *et al.* 2014; Sipkema *et al.* 2015; Miller *et al*. 2016; Russell *et al.* 2017; Russell and Cavanaugh 2017). This pattern is likely not explained by the taxa present, as the subsampled taxa analysis produced the same deficit of vertical transmission (Figure 3D) and a diversity of symbionts, from Gammaproteobacteria to Betaproteobacteria, have been shown to be strictly vertically transmitted on land. An internal symbiont transmission route does not appear to be a barrier to mixed transmission, as vertical transmission in vesicomyid and solemyid bivalves and sponges occurs through the female germline (Ikuta *et al.* 2016; Russell 2016) or brooded embryos (Schmitt *et al.* 2007), respectively, as in terrestrial associations. It should be noted, that the chemosynthetic flatworm *Paracatenula* may be strictly vertically transmitted, as it exhibits vertical transmission during asexual reproduction and patterns of host-symbiont co-speciation (Dirks *et al.* 2012), however, population-level data is needed to confirm this. It is interesting to consider that avoiding horizontal transmission in aquatic environments may be exceedingly difficult, even when cellular boundaries are present, and this may drive the lack of strict vertical transmission in these environments.

Explaining the abundance of strict horizontal transmission in the ocean is still problematic, as more information is needed to resolve the evolutionary pressures shaping these associations. There are two main possibilities for why strict horizontal modes persist in these taxa: 1) hosts and/or symbionts are incapable of germline integration or 2) it is not in the fitness interests of the symbiont or host to become vertically transmitted. In the case of the bobtail squid and its horizontally transmitted bioluminescent symbionts, horizontal transmission may be the optimal strategy for proper colonization timing. First, ample symbionts exist in the surrounding seawater for colonization of the light organ (Jones *et al*. 2006), indicating there is little danger of hosts not acquiring symbionts. Second, colonization is timed with host development and serves to induce formation of the organ itself (Nyholm and McFall-Ngai 2004). A similar situation exists for the hydrothermal vent tubeworm symbiosis, as symbionts are at high abundance in the surrounding seawater (Harmer *et al.* 2008) and colonize host tissues at a specific point in development, immediately after settlement and metamorphosis (Nussbaumer *et al.* 2006). In the case of the tubeworms, their habitat is patchy and the likelihood of any particular tubeworm offspring surviving to settlement and adulthood is low (Mullineaux *et al.* 2000), so symbionts may experience higher fitness by remaining somewhat independent of the host. Additionally, symbionts could be locally-adapted to their surroundings (Beinart *et al*. 2012), and so, hosts dispersing into the area and settling would be better served by picking up these bacteria compared to the bacteria housed in their parent’s tissues. So the only risks for these associations are not finding and acquiring symbionts in time for normal development, however some studies show that this may not be an issue and symbionts can be acquired by adults (Wentrup *et al.* 2014).

The enrichment for mixed mode transmission in both defensive and manipulative symbioses may be due to unique selection pressures acting on these associations relative to nutritional symbioses. In the case of defensive symbioses, the bacterial symbionts are often supplying an antiviral, antibiotic, or anti-parasitoid benefit, which likely requires exposure to these threats in the environment in order to respond to pathogen coevolutionary responses and maintain the defensive strategy, preventing the evolution of strict vertical transmission. Strict horizontal transmission may be problematic for associations which require bacterial protection during embryogenesis or larval stages (Kaltenpoth *et al.* 2006; Kölsch and Pedersen 2010). However, the success of the horizontally transmitted defensive symbiosis between *Aliivibrio fischeri* (formerly *Vibrio fischeri*), with its intricate signaling mechanisms for bacteria-host recognition and incorporation (Nyholm and McFall-Ngai 2004), indicates that horizontal transmission is permitted if symbionts are not required until adulthood. Although transmission mode data is lacking, chemical data suggests that marine defensive symbionts may be more prevalent than I have reported here (Flórez *et al.* 2015), and epibionts, which would likely be horizontally transmitted, are likely to be common (Lopanik 2014), so the horizontal transmission rate of defensive symbioses may be higher than presented here. In the case of manipulative associations, symbionts are adept at utilizing host resources for their own reproduction, precluding the evolution of a purely horizontal strategy, however, occasional horizontal transmission events increases their frequency in the host population, leading to the predominance of mixed modes. These strategies are distinct from those in nutritional symbioses, which experience pressures for host dependence (Fisher *et al.* 2017), such as metabolic complementarity between symbiont and host. The same is also likely true for multiple function symbioses, as nutrition is often one of the functions, and unknown symbioses, many of which are likely nutritional associations.

The non-random distribution of symbiont transmission modes among environment types, transmission routes, and functions has important implications for host and symbiont evolution. In regards to symbiont-specific impacts, the enrichment for horizontal/mixed modes in aquatic environments and vertical modes in terrestrial environments likely produces an excess of reduced symbiont genomes on land relative to in the ocean. While this is certainly anecdotally true (Moran and Bennett 2014), it was not tested here because genomes were not available for all of the included symbioses, but should be investigated in the future. How the rate of horizontal transmission impacts symbiont genome evolution remains one of the most important unanswered questions in symbiosis biology. In terms of host-symbiont interactions, the different distributions of transmission modes among symbiosis functions suggests that these groups may share suites of molecular characteristics. For example, defensive and manipulative symbioses are enriched in mixed mode transmission, thus they likely have specific cellular mechanisms for integration with host development and specific host recognition (Ebert 2013). Transmission mode also has important impacts on host evolution given that host tissues are often remodeled to accommodate symbionts and hosts must compensate for symbiont function or do without if symbionts are not acquired by the proper developmental stage (Kikuchi *et al.* 2011). Thus, hosts often exhibit morphological and developmental modifications relative to their ancestors that are tailored to symbiont biology, such as the light organ in the bobtail squid, *Euprymna scolopes* (Nyholm and McFall-Ngai 2004).

## Conclusions

The literature compiled in this study on bacterial symbiont transmission modes shows a strikingly non-random distribution between terrestrial and aquatic environments, transmission routes, functions, and among host/symbiont taxa. From the analysis presented above, it is reasonable to conclude that unless restricted by a host/symbiont function or an inhospitable external environment, bacterial symbionts tend to mix between hosts, which results in relatively high rates of mixed mode transmission across taxa (34.0%). However, rates are likely even higher, as a fair amount of data is needed to detect mixed modes (Figure S1, supporting information), so it often goes undetected. Regardless of the cause, there is clearly a bias for horizontal transmission in water and vertical transmission on land. Given that host restriction significantly reduces symbiont population size, driving genome degradation in strictly vertically transmitted associations, bouts of horizontal transmission may serve to increase the effective population size and mitigate some of the deleterious consequences of such obligate host-association, altering the evolutionary trajectories of symbioses in these different environments.

## Acknowledgements

I thank the Corbett-Detig, Cavanaugh, Haig, Sullivan laboratory members past and present, for helpful discussion and suggestions. I also thank the two reviewers for their insightful comments and suggestions, which helped improve the manuscript.

The author has no conflict of interest to declare.

